# Overexpression of Arabidopsis ubiquitin ligase AtPUB46 enhances tolerance to drought and oxidative stress

**DOI:** 10.1101/379859

**Authors:** Guy Adler, Amit Kumar Mishra, Tzofia Maymon, Dina Raveh, Dudy Bar-Zvi

## Abstract

The U-Box E3 ubiquitin ligase, AtPUB46, functions in the drought response: T-DNA insertion mutants of this single paralogous gene are hypersensitive to water- and oxidative stress (Adler et al. BMC Plant Biology 17:8, 2017). Here we analyze the phenotype of *AtPUB46* overexpressing (OE) plants. *AtPUB46*-OE show increased tolerance to water stress and have smaller leaf blades and reduced stomatal pore area and stomatal index compared with wild type (WT). Despite this, the rate of water loss from detached rosettes is similar in *AtPUB46-*OE and WT plants. Germination of *AtPUB46-*OE seeds was less sensitive to salt than WT whereas seedling greening was more sensitive. We observed a complex response to oxidative stress applied by different agents: *AtPUB46*-OE plants were hypersensitive to H_2_O_2_ but hyposensitive to methyl viologen. *At*PUB46-GFP fusion protein is cytoplasmic, however, in response to H_2_O_2_ a considerable proportion translocates to the nucleus. We conclude that the differential stress phenotype of the *AtPUB46*-OE does not result from its smaller leaf size but from a change in the activity of a stress pathway(s) regulated by a degradation substrate of the *At*PUB46 E3 and also from a reduction in stomatal pore size and index.

**Accession Numbers:** Sequence data for this article can be found in: The Arabidopsis Information Resource database (http://www.arabidopsis.org) under accession numbers At5G18320 (PUB46).

## 1. Introduction

Abiotic stress leads to a global change in plant proteostasis with the ubiquitin-26S proteasome system (UPS) being the key mechanism for regulated degradation of proteins [1-3]. Substrates are targeted for degradation by a pathway comprising ubiquitin-activating (E1), ubiquitin-conjugating (E2) enzymes and ubiquitin ligases (E3) [1-3]. Whereas E1 and E2 have a general role in protein ubiquitylation, E3s display high substrate specificity towards the target protein. Correspondingly, plants genomes contain small gene families encoding E1s and E2s, but high number of genes, e.g. over 1400 in Arabidopsis, that encode E3s [2]. Besides its role in regulated protein degradation ubiquitylation is also important in signaling and modification of cellular activities [3]. The UPS is involved in essentially all aspects of plant homeostasis such as growth and development, response to plant hormones and the response to abiotic and biotic stresses (reviewed by [3-6]). As sessile organisms that must respond to changes in their environment to survive, plants have the capacity to change their cellular homeostasis to better accommodate the changing environments, including stress conditions. These involves global changes in transcriptome, proteome, metabolome and iononome [7]. Regulated protein degradation directs the required specific changes in the cell proteome following exposure to abiotic stress conditions. Accordingly, a large number of E3s have been shown to be involved in the plant response to abiotic stresses in establishing altered proteostasis [4, 8-10].

We recently reported that the Arabidopsis E3, Plant U-box 46 (*At*PUB46), is involved in the response to abiotic stress [11]. The PUB family of monomeric U-box proteins comprise a ∼70 amino acid RING-like motif [8, 12, 13] followed by 3 non-identical repeats of the Armadillo (ARM) motif, shown to function in protein-protein interactions [14, 15]. *AtPUB46* is expressed in both roots and shoots [11]. *AtPUB46* transcript levels are modulated by abiotic stress: salt stress elevates its transcript levels in both roots and shoots, and osmotic stress enhances levels in the shoots. T-DNA insertion *Atpub46* mutants display hypersensitivity to both water and oxidative stress indicating a major role for *AtPUB46* in the abiotic stress response [11].

The marked pleiotropic stress response of *Atpub46* T-DNA insertion mutants led us to examine whether and how overexpressing *AtPUB46* would impact the stress response. Arabidopsis plants that overexpress *AtPUB46* from the strong cauliflower 35S virus promoter were constructed and tested for their response to those stresses in which absence of the protein led to hypersensitivity. Indeed we found that in contrast to the hypersensitive water-stress phenotype of the *Atpub46* T-DNA insertion mutants, *AtPUB46-*OE plants displayed enhanced water-stress survival. Given the contrasting phenotype attained by *AtPUB46-*OE we extended our study to salt and oxidative stresses applied by different agents. These experiments were conducted at different stages of plant development as previous reports have indicated that the stress reaction may be organ- and stage specific [4, 5, 8, 16]. *AtPUB46-*OE could exert its effect through leaf morphology and we therefore measured leaf size and stomatal index and whether this affected water loss in isolated leaf rosettes. The UPS is involved in gene regulation and we examined whether AtPUB46 protein is translocated into the nucleus in response to stress. An analysis of our results indicates a central role for the *AtPUB46-*encoded E3 in regulating the abiotic stress response.

## 2. Materials and methods

### 2.1. Plant material

#### *Arabidopsis thaliana* ecotype Columbia plants were studied

##### Plant transformation and selection of transgenic plants

*Agrobacterium* GV-3101 harboring the respective plasmids was used for genetic transformation of Arabidopsis by the floral dip method [17]. Transgenic seedlings were selected on plates containing 30 μg/ml hygromycin. Experiments were performed on T3 generation homozygous plants containing single-site T-DNA inserts. T-DNA insertion *Atpub46* mutants are described elsewhere [11].

### 2.2. Construct design

#### *AtPUB-OE* plants

Protein-encoding DNA sequences were amplified by PCR using genomic DNA as a template and gene specific primers (Table S1). Amplified DNA fragments were sub-cloned into the plant transformation vector pCAMBIA99-1 downstream of the constitutive strong *CaMV 35S* promoter.

#### Constructs for expression of *At*PUB46-eGFP fusion proteins

cDNA amplified fragments were fused to the N-terminus of eGFP in the pSAT4-eGFP-N1 plasmid [18] downstream of the constitutive *CaMV 35S* promoter. The *CaMV 35S::AtPUB46-eGFP* fusion cassette was ligated into pCAMBIA 1302 replacing the *CAMV 35S::6xHIS-GFP* sequence of the vector.

### 2.3. Transcript levels

RNA isolation, cDNA synthesis and RT-qPCR assay for determining relative steady state transcript levels were performed as previously described [19]. Primers are listed in Table S1.

### 2.4. Plant growth and stress application

Surface sterilized seeds of the indicated genotypes were cold treated and sown in Petri dishes containing half strength Murashige and Skoog (0.5 x MS) nutrient solution mix [20], 0.5% sucrose and 0.6% agarose, or in pots containing planting mix as previously described [21]. Plants were grown at 22-25 °C, 50% humidity under continuous light or a 12 hr light/dark regime. Where indicated, plates contained in addition the indicated concentrations of ABA, hygromycin, or abiotic-stress agents (NaCl, mannitol, H_2_O_2_ or methyl viologen (MV)).

#### 2.4.1. Drought tolerance

*AtPUB46-*OE and control WT plants were grown

for 3 weeks in pots with equal amounts of potting mix under non-stressed conditions. Water was then withheld and plant wilting was followed daily for a further 2 weeks.

#### 2.4.2. Seed germination and cotyledon greening assay

Surface-sterilized cold-treated seeds were sown on Petri plates containing 0.5 x MS, 0.7% agar and where indicated NaCl, mannitol, MV or ABA. Seed germination (root emergence) was scored 2 days after sowing for NaCl, mannitol and H_2_O_2_ treatments, and after 6 days for ABA treatment. Seedling greening was assayed 5 days after sowing for MV treatment, and 12 days after sowing for all other treatments.

### 2.5. Size of cotyledon and analysis of leaf epidermal cells

#### 2.5.1. Cotyledon surface area

50 cotyledons were excised from 5 day old seedlings, and placed on glass plates together with a 25 mm^2^ calibration scale. Plates were scanned using a HP Scanjet 5400C at 600 dpi.

#### 2.5.2. Epidermal cell size and number

The 3rd and 4th leaves of 5-week-old plants were painted with transparent nail polish. After drying, the painted layer was peeled off, and photographed using a light microscope. The number of stomata and pavement cells was determined using the cell counter plugin of ImageJ software as were the area of cotyledons and leaf epidermal cells. Stomatal and pavement cell density (number in mm^2^) was calculated. Stomatal index [Stomata * 100/(Stomata + pavement cells)] was calculated according to Royer [22].

#### 2.5.3. Stomatal aperture area

3rd and 4th leaves of 4-week-old plants were treated with stomata-opening buffer as previously described [23]. The lower epidermis was peeled off and the stomatal aperture area measured using the NIS-Elements D 4.2 software (Nikon, Japan).

### 2.6. Water loss

Rosettes of one-month old plants were cut and placed on weigh boats with their abaxial side up. Rosette weight was determined immediately after cutting and at 10 min intervals. Weight of each rosette at different times after detachment was normalized to its weight at time zero.

### 2.7. Photosynthetic efficiency

Photosynthetic efficiency of photosystem II was assayed in leaves of pot-grown plants using a MINI-PAM-II fluorometer (Walz GmbH, Effeltrich, Germany).

Photosynthetic efficiency, presented as Fv/Fm was measured and calculated as previously described [11].

### 2.8. Subcellular localization

Two-week old plate grown seedlings of transgenic plants expressing *35S::AtPUB46-eGFP* constructs were transferred to plates containing Whatman No. 1 filter paper soaked with 0.5 x MS, 0.5% sucrose, or with this medium containing in addition 200 mM NaCl, 400 mM mannitol, 1 μM MV, or 100 mM H_2_O_2_. Plates were incubated for 6 h in the light, and roots were examined with a Zeiss LSM880 confocal microscope. Where indicated, roots of the harvested seedlings were also incubated for 5 min in DNA staining solution containing 1 mg/ml DAPI in 0.1% triton X-100.

### 2.9. Statistical analysis

Each experiment was performed at least three times with over 50 plants for each treatment. Results are presented as mean ± SE [calculated using SPSS software version 18, (SPSS Inc, Chicago, IL)]. Differences between groups were analyzed by Tukey’s HSD post-hoc test (P≥0.05).

## 3. Results

### 3.1. *AtPUB46-*OE plants display enhanced tolerance to drought stress Arabidopsis plants were transformed with pCAMBIA encoding *CaMV*

*35S::AtPUB46*. Lines resulting from single T-DNA insertion events were selected, and seeds of the respective homozygous plants were collected. Steady-state transcript levels of *AtPUB46* in all transgenic lines were a thousand fold higher than the levels of the endogenous gene transcripts in WT seedlings (Fig. S1).

T-DNA insertion *Atpub46* mutants are hypersensitive to drought stress compared with WT plants [11]. We therefore challenged pot-grown *AtPUB46*-OE WT plants to 2 weeks of water withholding. *AtPUB46*-OE resulted in improved tolerance to drought stress (Fig. 1A-B). These results were essentially independent of the rosette size; data obtained from comparing drought sensitivity of *AtPUB46*-OE and WT plants of similar age (and different size), or similar size (and different age) were indistinguishable from those obtained from comparing same age plants. In contrast to WT plants in which the photosynthetic apparatus was hypersensitive to water withholding, in *AtPUB46*-OE plants the photosynthetic efficiency of irrigated plants was unaffected throughout the experiment irrespective of the duration of water withholding (Fig. 1C).

**Figure 1.**
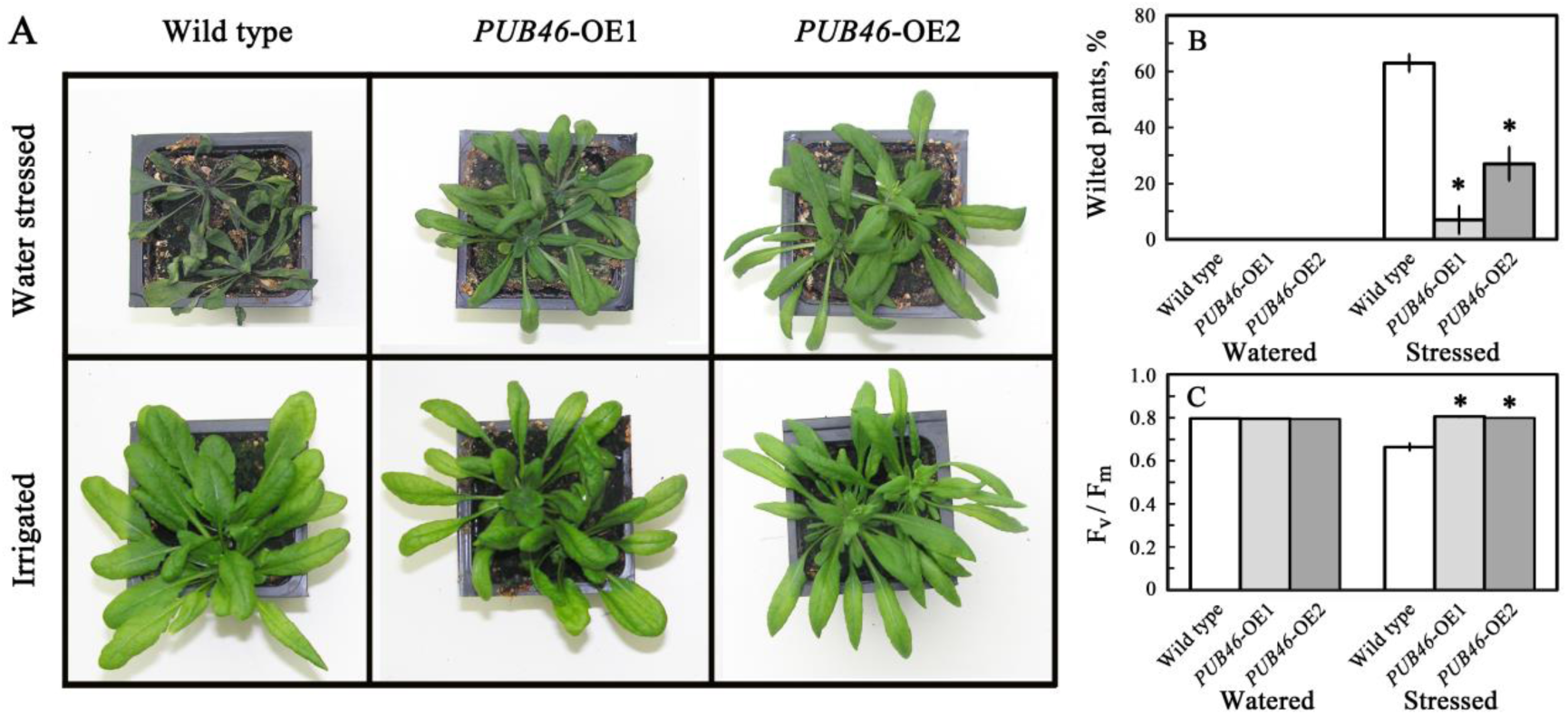
Water stress performance of pot-grown *AtPUB46-OE* plants. Potted plants were grown for one month followed by water withholding for 2 weeks. (A) Photos of representative plants. (B) Quantitative data of wilted plants. (C) Photosynthetic efficiency. All data shown are average ± SE. Statistically significant changes from WT plants in the same treatment (P<0.05) are marked with an asterisk.

### 3.2. Overexpressing *AtPUB46* affects leaf and stomata morphology

*AtPUB46*-OE plants showed reduced leaf blade width but no difference in leaf length or the number of leaves in the rosette (Fig. 2A-C). The reduced leaf area results from a reduction in cell size rather than cell number (Fig. 2D-F). This was assayed by measurement of the size and density of pavement cells of the lower epidermis: these cells are significantly smaller in *AtPUB46*-OE plants than in WT plants (Fig. 2D-F) and cell density is correspondingly higher (Fig. 2G). Interestingly, cotyledons of the *AtPUB46*-OE seedlings were also smaller than those of WT seedlings (Fig. S2).

**Figure 2.**
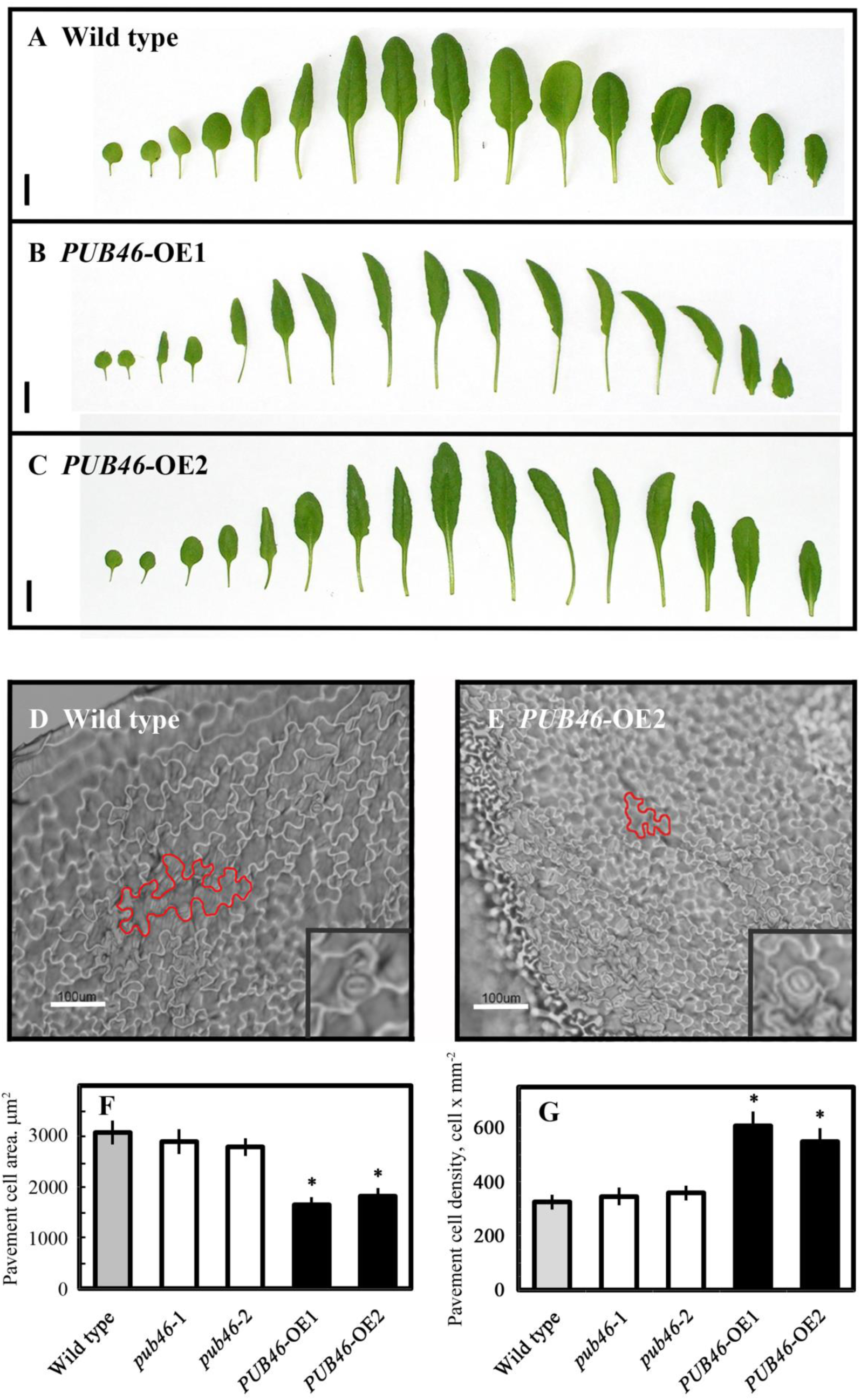
*AtPUB46*-OE have reduced leaf size. One month old pot-grown plants were analyzed for the following parameters: (A-C) dissected rosette leaves; (D-G) lower epidermis of the 5^th^ leaves of WT and *AtPUB46*-OE: Lower epidermis cell print of WT (D) and *AtPUB46-*OE (E) plants, borders of representative pavement cells are marked with a red line, Bar=100 μm, inserts show stomata at 2 x magnification. Cell area (F) and cell density (G) of pavement cells in leaves of WT, *Atpub46*-T-DNA insertion mutants and *AtPUB46*-OE plants. Data shown are average ± SE. Statistically significant changes from WT plants (P<0.05) are marked with an asterisk.

Changes in drought tolerance may result from altered water loss from the leaves. However, water loss from detached rosettes of WT and *AtPUB46*-OE was not significantly different (Fig. 3A), indicating that transpiration rates of these plants may be similar and not the reason for the enhanced drought tolerance of the *AtPUB46*-OE plants. We therefore looked in greater detail at the stomata of the lower epidermis of the rosette leaves. Overall, stomata of *AtPUB46*-OE plants were smaller than those of WT plants and *Atpub46*-T-DNA insertion mutants (Fig. 3B). Whereas stomatal density was similar in all the studied plant genotypes (Fig. 3C), the stomatal index was significantly lower in leaves of *AtPUB46*-OE plants than in WT and T-DNA insertion mutants (Fig. 3D). Stomatal pore area was measured both in open and in induced-closed states. When fully open, the stomatal pore area of *AtPUB46*-OE plants is smaller than that of WT and T-DNA insertion mutants. Stomata of all tested lines were responsive to ABA. In the ABA-induced closed state, the stomatal pore area of the *AtPUB46*-OE plants was similar to that of WT plant, whereas the pore area of the *Atpub46*-T-DNA insertion mutants was larger (Fig. 3E).

**Figure 3.**
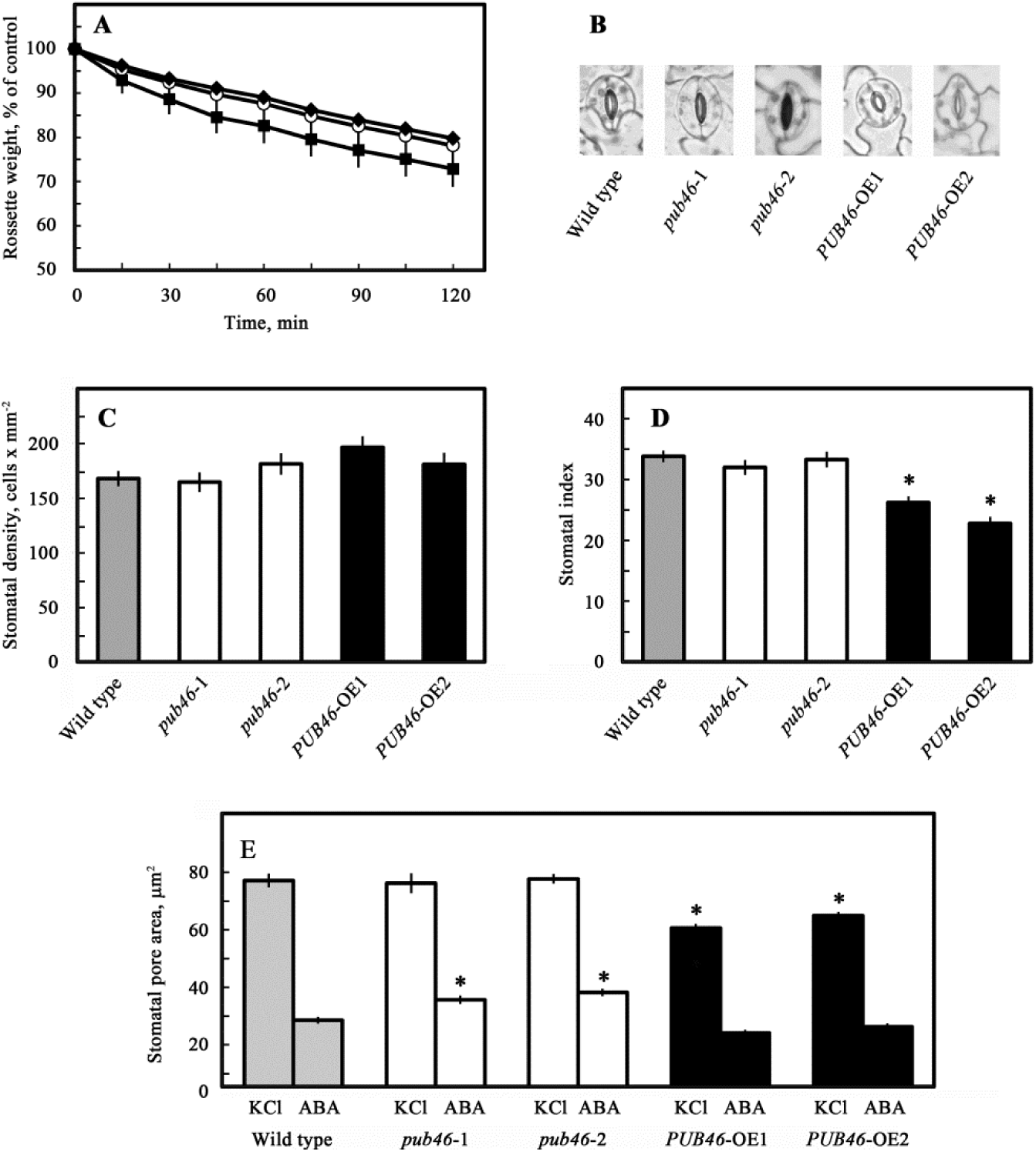
Overexpression of *AtPUB46* affects stomatal index and pore area but not water loss rate. (A) Water loss. Rosettes of WT (white circles), AtPUB46-OE1 (Black diamond), or AtPUB46-OE2 (black square) were detached and weighed at the indicated times. Changes obtained for the three genotypes were not statistically significant. (B) Pictures of representative stomata of WT, *Atpub46* T-DNA insertion mutant and *AtPUB46*-OE. Stomatal density (C) and index (D) in lower epidermis of full size rosette leaves. (E) Stomatal pore area of leaves treated with KCl or ABA. Data shown are average ± SE. Statistically significant changes from WT plants in the same treatment (P<0.05) are marked with an asterisk.

### 3.3. Overexpressing *AtPUB46* affects seed germination and greening

We assayed germination of *AtPUB46*-OE under abiotic stress conditions. Under control conditions, germination of seeds was indistinguishable from that of WT seeds, reaching essentially 100% germination (Fig. 4). However, under stress conditions there was a clear difference between control and OE plants: WT seed germination in 150 mM NaCl was reduced by ca. 50% whereas germination of the *AtPUB46*-OE seeds was only slightly affected (Fig. 4A). In contrast a similar degree of inhibition of germination was observed in WT and *AtPUB46*-OE plants in response to osmotic stress administrated by 300 mM mannitol (Fig. 4B), and to hormone treatment with 1 μM ABA (Fig. 4C). Seed germination of *AtPUB46*-OE plants was more sensitive to 5 mM H_2_O_2_ than WT (Fig. 4D). Unlike its differential effect on root emergence, NaCl inhibition of root development was similar in WT and *AtPUB46-*OE plants (Fig. S3).

**Figure 4.**
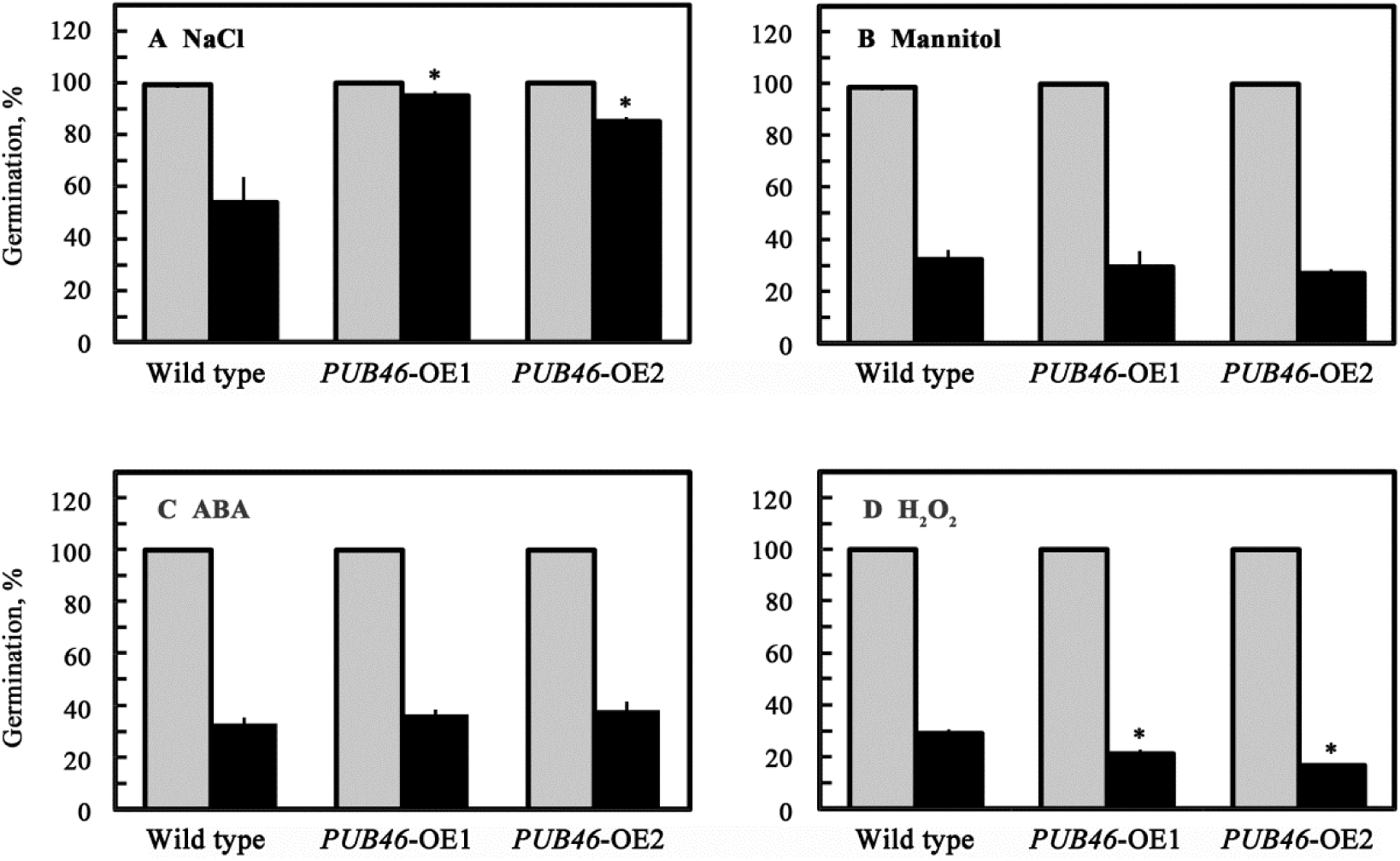
Effects of overexpressing *AtPUB46* on seed germination. Surface sterilized cold treated seeds of the indicated plant lines were plated on agar solidified media containing 0.5 x MS, 0.5% sucrose (control) supplemented with: (A) 150 mM NaCl; (B) 300 mM mannitol; (C) 1 μM ABA or (D) 5 mM H_2_O_2_. Germination scored 2 and 6 days later for (A, B, D) and (C), respectively. Gray bars, non-treated; black bars, treated seedlings. Data shown are average ± SE. Statistically significant changes from WT plants in the same treatment (P<0.05) are marked with asterisks.

We also determined the effect of different stress agents on seedling greening of WT and *AtPUB46*-OE. Under normal conditions, 100% of the seedlings of all tested genotypes are green (Fig. 5 & Fig. S4). In the presence of 150 mM NaCl, ca. 50% WT seedlings remained green 12 days after sowing, whereas almost all *AtPUB46*-OE seedlings were bleached by the same treatment (Fig. 5A). A similar percent of green seedlings in WT and *AtPUB46*-OE was observed in response to osmotic stress with 300 mM mannitol, and in response to 1 μM ABA (Fig. 5B & C). 5 mM H_2_O_2_ reduced the greening of WT and *AtPUB46*-OE seedlings with the *AtPUB46*-OE seedlings being more sensitive than WT (Fig. 5D). In contrast, a higher percentage of the *AtPUB46*-OE seedlings remained green when exposed to oxidative stress by the photosynthesis-dependent agent MV (Fig. 5E).

**Figure 5.**
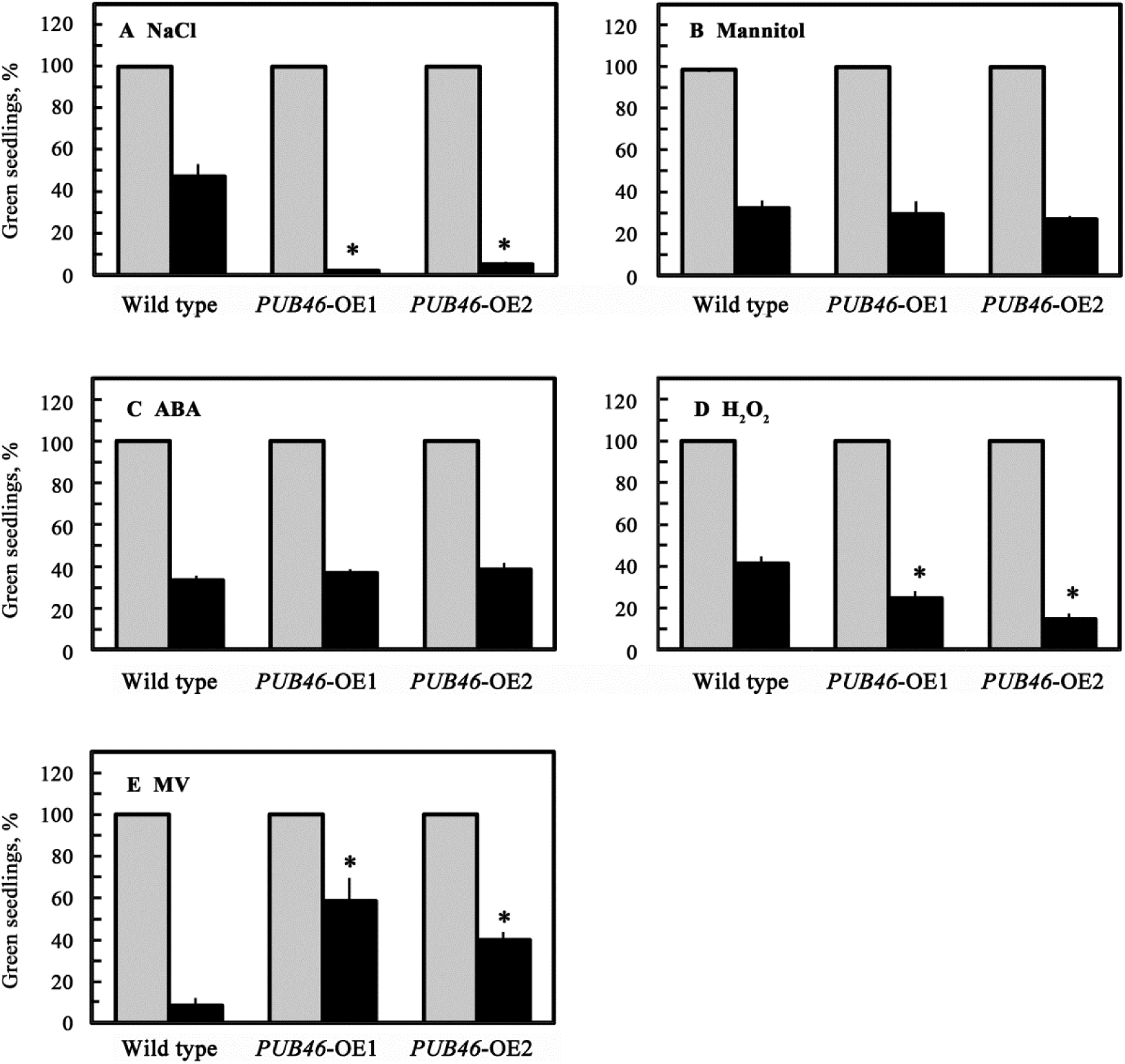
Effects of overexpressing *AtPUB46* on seedling greening. Surface sterilized cold treated seeds of the indicated plant lines were plated on agar plates with 0.5 x MS, 0.5% sucrose (control) supplemented with: (A) 150 mM NaCl; (B) 300 mM mannitol: (C) 1 μM ABA; (D) 5 mM H_2_O_2_; (E) 1 μM methyl viologen (MV). Green seedlings were scored 12 (A-D) or 5 (E) days later. Gray bars, non-treated; black bars, treated seedlings. Data shown are average ± SE. Statistically significant changes from WT plants in the same treatment (P<0.05) are marked with asterisks. Representative plates are shown in Fig. S4.

### 3.4. *At*PUB46 subcellular localization

To determine the subcellular distribution of *At*PUB46 we transformed Arabidopsis with pCAMBIA-*CaMV 35S::AtPUB46-eGFP*. The fluorescent *At*PUB46-eGFP protein was detected in the cytosol (Fig. 6A). Treatment with the stress agents NaCl, mannitol and MV did not affect the strength or localization of the fluorescent signal (Fig. 6B-D). In contrast, roots of plants treated with 100 mM H_2_O_2_ showed an additional strong nuclear fluorescent signal (Fig. 6E, arrows). Nuclear localization in response to H_2_O_2_ treatment was confirmed by co-localization of the *At*PUB46-eGFP fluorescent protein with the DAPI DNA binding probe (Fig. 6F-H). In contrast, H_2_O_2_ treatment did not significantly affect the cellular localization of the eGFP tag, when expressed alone (Fig. S5), suggesting that the translocation to the nuclei was AtPUB46-dependent.

**Figure 6.**
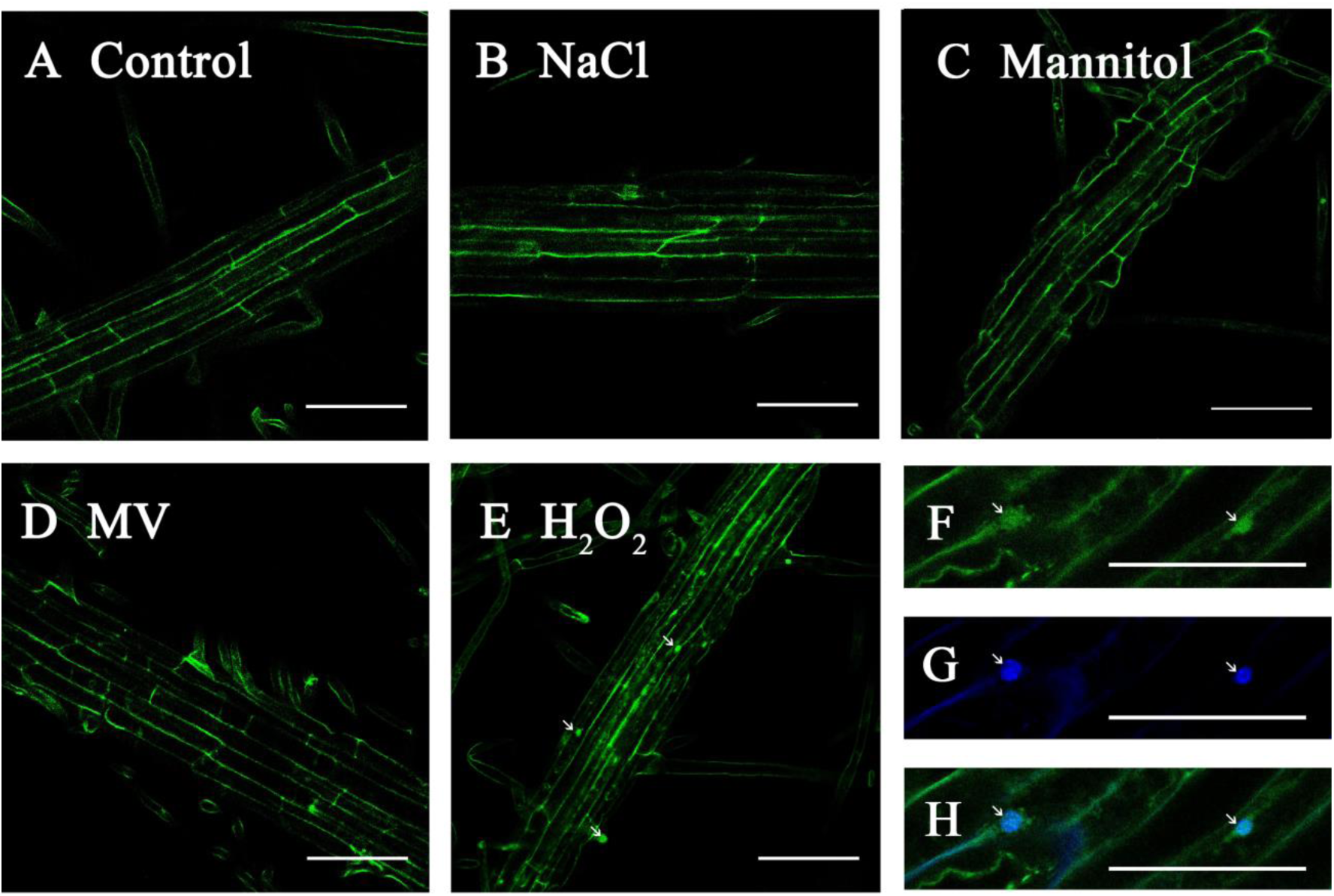
Subcellular localization of AtPUB46. Roots of 2 week old transgenic Arabidopsis plants expressing AtPUB46-eGFP fusion proteins driven by the CaMV 35S promoter were transferred to a Petri dish with filter paper soaked with 0.5 x MS salts, 0.5% sucrose (A), or this medium complemented with 200 mM NaCl (B); 400 mM mannitol (C); 1 μM methyl viologen (MV) (D); 100 mM H_2_O_2_. (E-H): Plates were incubated for 6 h under illumination. F-H, roots were incubated with DAPI. Roots examined by confocal fluorescent microscopy set for eGFP (A-F) or DAPI (G) fluorescence. Panel H show merged images F and G. Bars = 50 μm.

## 4. Discussion

### 4.1. *AtPUB46* is involved in the plant response to abiotic stress

T-DNA insertion *Atpub46* mutants are hypersensitive to drought stress and water withholding affects their photosynthesis efficiency compared with WT plants [11]. These findings indicate a major role for *AtPUB46* in plant drought tolerance [11]. Here we show that overexpressing *AtPUB46* plants leads to enhanced tolerance to drought both with respect to wilting and photosynthesis under water stress. These results extend and confirm the role of AtPUB46 E3 in the response to water stress.

Monomeric E3s of the PUB-family are frequently involved in the plant stress response both as positive or negative regulators [4, 5, 8-10, 16, 24-27]. Our results indicate that *AtPUB46* is a positive regulator of the response to drought as OE plants display improved tolerance and knock-down mutants are hypersensitive [11]. This is similar to OE of rice *OsPUB15* [28], or wheat *TaPUB1* [29] in transgenic rice or *Nicotiana benthamiana,* respectively, which also increased drought tolerance. This is in marked contrast to the majority of PUB genes that have been studied where overexpression resulted in hypersensitivity to drought [30-35].

Overexpressing *AtPUB46* had differential effects on the plant response to different abiotic stress agents: seeds of *AtPUB46-*OE plants showed better germination than WT in the presence of 150 mM NaCl. In contrast, salt induced a higher extent of bleaching in the developed seedlings. Similar results were reported in durum wheat where salt and osmotic stresses affected germination and later developmental stages differently [36]. Moreover, the capacity of three Arabidopsis *RS* mutants (*RS17, RS19* and RS20) to tolerate NaCl was shown to be dependent on the developmental stage, differences being observed only at germination [37]. The differential effect on inhibition of germination compared with seedling greening between WT and *AtPUB46*-OE can be attributed to the toxic effect of the salt, rather than to its osmotic component, since a similar extent of osmotic stress, administrated by with 300 mM mannitol instead of 150 mM NaCl, did inhibit germination and greening differentially. Furthermore, ABA, the plant hormone related to abiotic stresses such as water-, osmotic- and salt-stresses (for a recent review see [38]) did not affect the sensitivity to inhibition of germination and greening of *AtPUB46*-OE plants, indicating that the NaCl response is not mediated by osmotic stress or ABA. Interestingly, ABA also does not affect the steady state levels of *AtPUB46-*transcripts [11].

Furthermore, whereas overexpressing *AtPUB46* dramatically decreased seedling greening in the presence of NaCl, knocking-down *PUB46* did not alter the fraction of green seedlings following the same treatment [11], suggesting that seedling greening in the presence of NaCl is not simply affected by the protein levels of AtPUB46, but has a more complex regulation.

Our data suggest that although salt-, osmotic-, and oxidative stress share some common components (for example, salt stress has an osmotic component, and both salt and water stress induce oxidative stress [39]), the mechanisms of the plant response to salt-, osmotic-, and oxidative stress are nevertheless distinct. Differences in the physiological responses to salt- and water stress have been attributed to genes whose activities control the transport of salt ions across membranes; these salt-specific effects may impact later growth more than germination [40]. Moreover, transcriptome and proteome analyses of plants exposed to drought- or salt stress, revealed that despite significant overlap in the pattern of gene and protein expression, each stress also affected a specific set of genes or proteins, suggesting that the molecular impact of these stresses is not redundant [41-47].

Furthermore, ROS administrated by the addition of H_2_O_2_ or MV (inducer of O_2_^-^) had different effects on the *AtPUB46*-OE plants. Although different ROS may react with and oxidize biological molecules such as lipids, carbohydrates, nucleic acids and proteins [48], ROS agents differ in their reactivity level, life span and their capacity to translocate within the cell or even between cells. MV is known to mediate electron transfer to molecular oxygen at the acceptor site of PSI [49]. The resulting short lived O_2_^-^ may react with molecules in its vicinity which can be hydrogenated to yield highly reactive hydroxyl radicals, or converted by superoxide dismutase to H_2_O_2_ [50]. H_2_O_2_ is a less reactive ROS that may even diffuse across membranes via aquaporin channels. As a result, H_2_O_2_, as well as other ROS also act in signaling in plants [50].

### 4.2. Drought tolerance of *AtPUB46*-OE may result from reduced stomatal pore area and index

The increased drought tolerance of plants with small leaves is often due to reduced transpiration from the smaller leaf area and by redirection of energy to processes involved in drought tolerance [51, 52]. Almost all transpiration occurs via the stomata, and conductance through stomata is modulated by their aperture and density [53]. Although leaves of *AtPUB46*-OE are smaller than those of WT plants, their stomatal density is unaltered. Stomata of the *AtPUB46-*OE plants have reduced aperture compared with WT plants and rates of water loss from rosettes of *AtPUB46-*OE plants were not significantly different from those of WT rosettes. When we compared plants of similar rosette size (achieved by comparing *AtPUB46*-OE plants with younger WT plants) we observed improved drought tolerance of the *AtPUB46*-OE plants. Thus, leaf size in itself is probably not the direct cause of the reduced transpiration.

Increased drought tolerance was observed in tomato plants that overexpressed the transcription factor SlDREB [54]. *SlDREB*-OE plants have suppressed GA biosynthesis and leaf expansion, reduced stomatal density and enhanced drought resistance. Overexpressing Arabidopsis *GIBBERELLIN METHYL TRANSFERASE 1* (*AtGAMT1*) in tomato increased drought tolerance [55]. Overexpressing *AtGAMT1* inhibited the expansion of leaf epidermal cells, leading to the formation of smaller guard cells resulting in smaller stomata with reduced stomatal pores, and corresponding reduced stomatal conductance [55]. We therefore argue that the impact of overexpressing *AtPUB46* on drought tolerance results primarily from a decrease in stomatal pore area and index, and not from reduced leaf size.

### 4.3. *At*PUB46 subcellular localization

The AtPUB46-eGFP fusion protein was detected in the cytoplasm. This subcellular localization is in agreement with subcellular prediction algorithms [56] and suggests that protein targets of PUB46 may consist primarily of cytoplasmic proteins. Treatment of seedlings with H_2_O_2_, but not with NaCl, mannitol or MV, resulted in localization of AtPUB46 in the nucleus in addition to the cytosol. Analysis of the amino acid sequence of PUB46 for putative NLS, reveals that it does not appear to have a “strong” NLS. It has two ‘suboptimal’ putative NLSs suggested by the cNLS Mapper and seqNLS algorithms [57, 58]. Interestingly, the score awarded to these sequences by the cNLS Mapper software is “localized to both nucleus and cytoplasm”. Indeed, this is what is expected of a modulated-localization signal which allows the protein localization to be modified. Proteasomes are present in both the cytoplasm and the nucleus [1-3]. Proteasome subunits are synthesized in stoichiometric amounts suggesting that transcription of their genes is coordinately regulated. Under a multitude of mild stress conditions proteasome levels rapidly rise concomitant with the need for removal of irreversibly damaged proteins and they may relocalize in response to localized concentrations of degradation substrates [59]. Thus, our finding suggests that although AtPUB46 appears within the cytosol, it translocates to the nucleus in response to specific signals, such as H_2_O_2_, implying the likelihood of nuclear degradation substrates. Subcellular localization of *At*PUB44 was altered by treatment with the plant hormones auxin and ABA, but was not affected by ethylene and gibberellin [27]. Similarly the COP1 E3 moves between the cytosol and the nucleus where it appears to function [45].

### 4.4. Concluding remarks

In a previous study, we showed that although *At*PUB46, *At*PUB47 and *At*PUB48 are very similar proteins encoded by paralogous genes, *At*PUB46 has an unique function in the response to water stress as single homozygous mutants of *AtPUB46* are hypersensitive to water stress [11]. Here we extend our analysis of the role of this E3 in the stress response with plants that overexpress *AtPUB46. AtPUB46*-OE plants display enhanced drought tolerance, and a complex response towards salt and oxidative stresses. Although overexpressing *AtPUB46* results in reduced leaf and stomata pore size and index, these structural changes do not suffice to explain the increased drought tolerance of the *AtPUB46-*OE plants. We therefore attribute the differential response to the various abiotic stresses examined here to an effect on cellular proteostasis. The *At*PUB46 target protein(s) whose degradation enhances plant tolerance to drought and oxidative stress remain to be identified.

## Acknowledgments

This work is supported by a grant from the Israel Science Foundation (to DR and DBZ) and by the I-CORE Program of the Planning and Budgeting Committee and the Israel Science Foundation (Center No. 757, to DBZ). DBZ is the incumbent of The Israel and Bernard Nichunsky Chair in Desert Agriculture, Ben-Gurion University of the Negev.

## Supporting Information

### Appendix A – Figures S1-S5

**Figure S1.** Relative expression of the *AtPUB46* in the respective overexpressing plants.

**Figure S2**. Area of cotyledons of *Atpub46* T-DNA insertion mutants and *AtPUB46-*OE plants.

Figure S3. Effect of NaCl on root length.

Figure S4. Effects of overexpressing *AtPUB46* on seedling greening.

Fig. S5. Effect of H_2_O_2_ treatment on the cellular localization of eGFP.

### Appendix A – Table S1

**Table S1.** List of primers used in this study.

---

**Figure S1.**
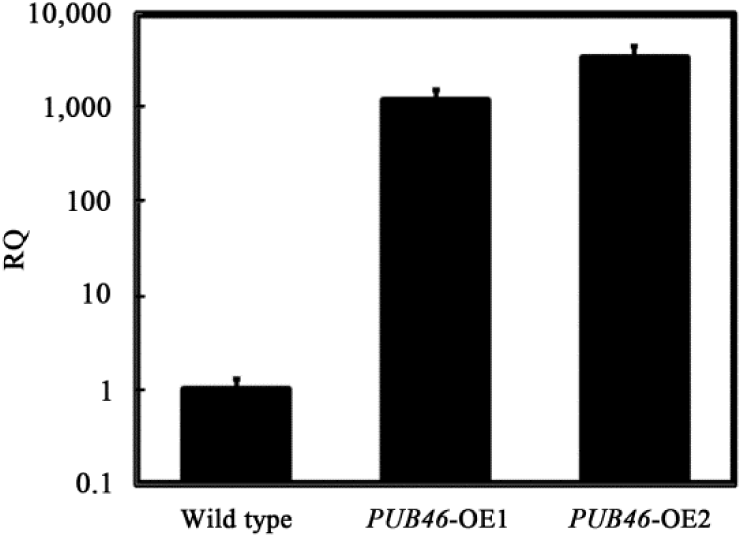
Relative expression of the *AtPUB46* in the respective overexpressor plants. mRNA was isolated from two week old WT and *AtPUB46*-OE seedlings. cDNA was prepared and analyzed by qPCR as described in methods. Transcript levels of *AtPUB46* in WT plants was defined as 1.

**Figure S2.**
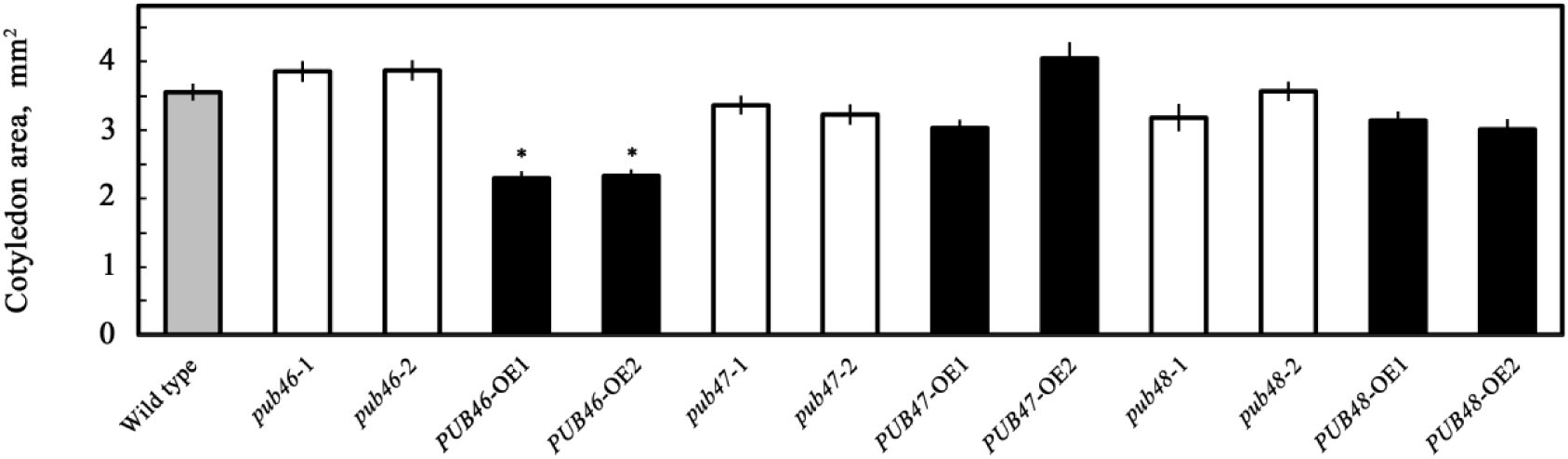
Area of cotyledons of *Atpub46* T-DNA insertion mutants and *AtPUB46-*OE plants. Area of cotyledons of four days old seedlings of the indicated lines was determined. Statistically significant changes from WT plants in the same treatment (P<0.05) are marked with asterisks.

**Figure S3.**
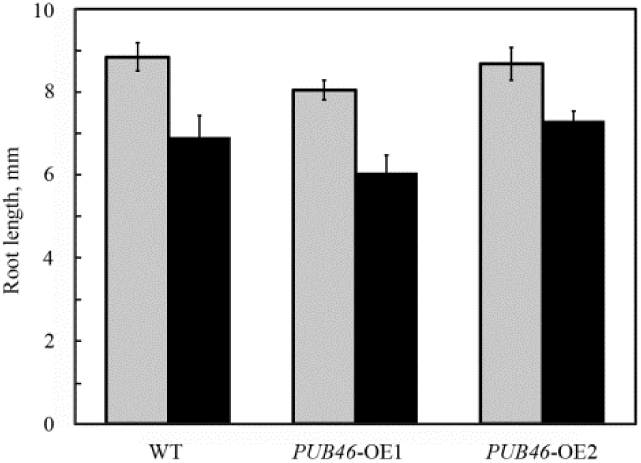
Effect of NaCl on root length. Surface sterilized cold treated seeds of the indicated plant lines were plated on agar plates with 0.5 x MS, 0.5% sucrose (gray bars) or on plates with this medium supplemented with 150 mM NaCl (black bars). Root length was measured 12 days later. Data shown are average ± SE.

**Figure S4.**
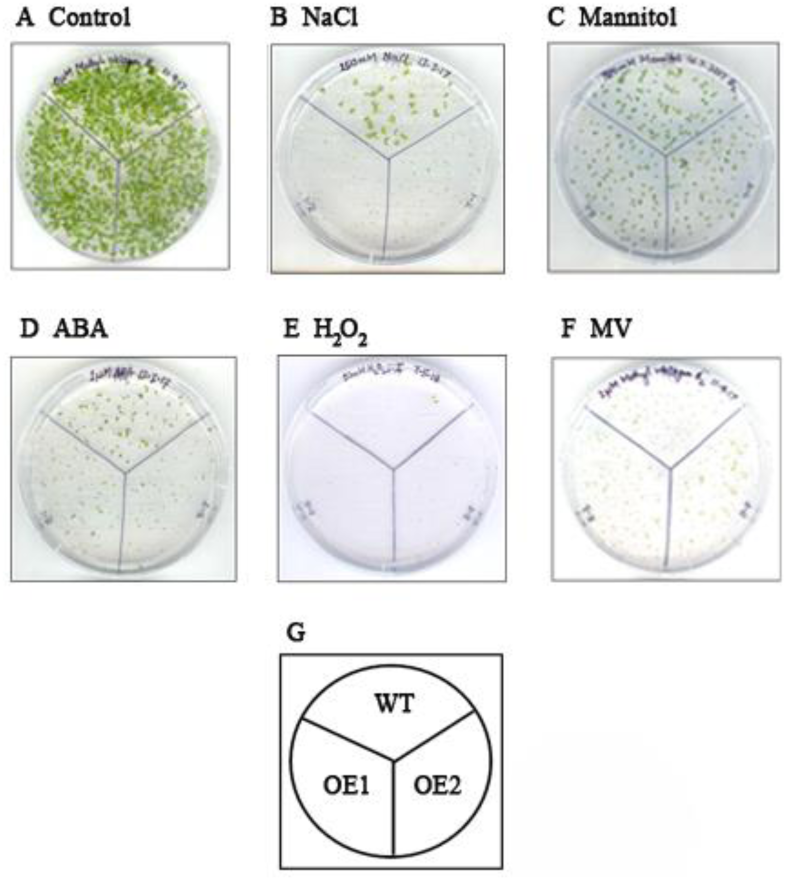
Effects of overexpressing *AtPUB46* on seedling greening. Surface sterilized cold treated seeds of the indicated plant lines were plated on agar plates with 0.5 x MS, 0.5% sucrose (control, A), or in the same medium supplemented with: (B) 150 mM NaCl; (C) 300 mM mannitol: (D) 1 μM ABA; (E) 5 mM H_2_O_2_; (F) 1 μM methyl viologen (MV). Green seedlings were scored 12 days later. Representative plates are shown in this figure. Quantitative analyses of the results are shown in Fig. 5.

**Figure S5.**
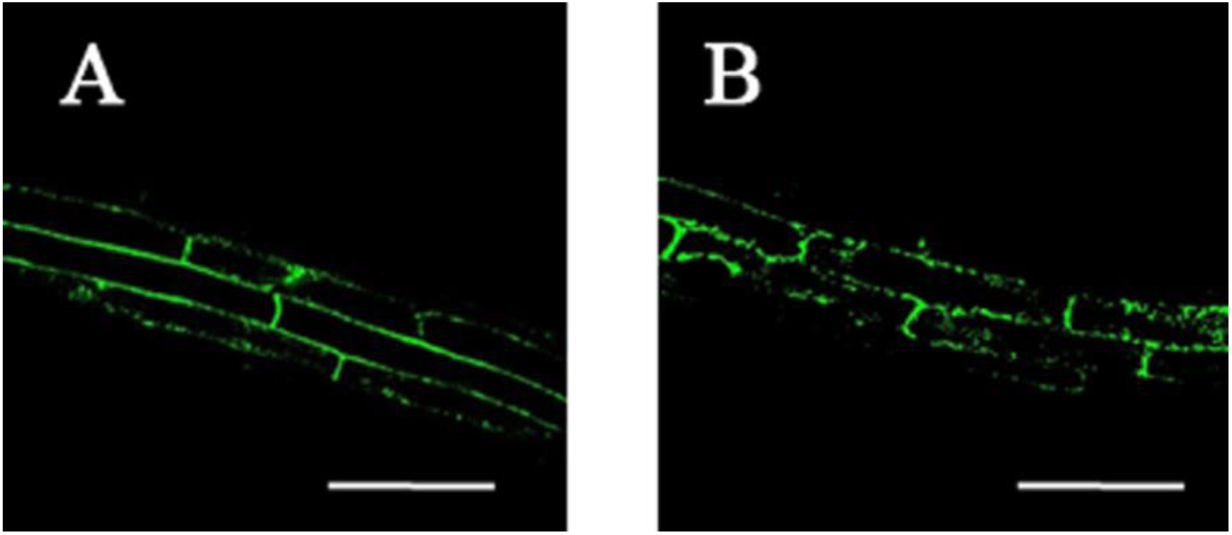
Effect of H_2_O_2_ treatment on the cellular localization of eGFP. One week old transgenic Arabidopsis plants expressing eGFP driven by the CaMV 35S promoter were transferred to a Petri dish with filter paper soaked in 0.5 x MS salts, 0.5% sucrose (A), or this medium complemented with 100 mM H_2_O_2_. (B). Plates were incubated for 6 h under illumination. Roots were examined by confocal fluorescent microscopy set for eGFP fluorescence. Bars = 50 μm.

